# Dissociation between impaired explicit spatial remapping and preserved implicit neural dynamics in Alzheimer’s disease

**DOI:** 10.64898/2026.07.06.736718

**Authors:** Xueling Wang, Yimeng Wang, Keliang Pang, Chenguang Zheng

## Abstract

The hippocampus supports spatial memory by dynamically integrating external sensory inputs with intrinsic neural circuit dynamics during novel experience. In Alzheimer’s disease (AD), despite impaired hippocampal spatial remapping, learning and memory-related hippocampal representations can remain partially preserved, a phenomenon consistent with cognitive resilience (Gómez-Isla and Frosch 2022, Jia, Xu et al. 2025). However, the hippocampal ensemble coding patterns associated with these preserved memory abilities remain remains unclear. We hypothesize that intrinsic temporal structures of neuronal firing continue to facilitate the encoding of new spatial information. Using the App^NL-G-F^ rat model, we recorded hippocampal CA1 activity across multiple days during a familiar-novel context alternating task. We found a dissociation between impaired explicit spatial coding and preserved implicit temporal coding in the AD hippocampal network. Explicit spatial coding was impaired, as place cells showed weak discrimination between distinct contexts and failed to exhibit improvement across days. In contrast, implicit temporal coding exhibited experience-dependent refinement, with cofiring dynamic becoming increasingly context-specific across long-term experience. Further analysis suggested that the enhancement of implicit cofiring may be associated with the increased consistency of neural ensemble reactivation during sharp wave ripples in awake rest. Taken together, these findings reveal an explicit–implicit dissociation in the AD hippocampal network, suggesting that the experience-dependent refinement of implicit temporal coding may support preserved learning-related hippocampal network dynamics despite impaired spatial remapping.

## Introduction

A fundamental computational function of the brain is to dynamically integrate fluctuating external sensory inputs with the intrinsic dynamics of neural circuits, thereby constructing coherent cognitive representations. As a pivotal hub for this integration, the hippocampus not only encodes external spatial locations but also organizes information through the temporal structure of intrinsic population activity (Rubin, Sheintuch et al. 2019, Nieh, Schottdorf et al. 2021, Levy, Carrillo-Segura et al. 2023, Nakai, Kitanishi et al. 2024). These complementary forms of representation can be described as explicit spatial coding, in which neuronal firing directly reflects external spatial location, and implicit temporal coding, which is expressed through the temporal relationships among neurons. They that may rely on distinct strategies but jointly support memory encoding. As animals explore distinct contexts, the hippocampal network rapidly registers the internal neural manifold to specific external spatial contexts, a process traditionally termed “remapping” (Fenton 2024). This mechanism enables the hippocampus to utilize limited neural resources to rapidly discriminate between similar or distinct spatial contexts (Dong, Madar et al. 2021) by adjusting the alignment between internal activity patterns and external cues. Subsequently, with the accumulation of experience, this registration is further stabilized via synaptic plasticity, leading to the gradual refinement and stabilization of spatial representations, which serves as the neural basis for robust long-term episodic memory (Pettit, Yap et al. 2022, Vaidya, Li et al. 2025).

In Alzheimer’s disease (AD), hippocampal place cell remapping is impaired during context transitions, reflecting reduced neural discrimination between distinct spatial contexts (Jun, Bramian et al. 2020). Furthermore, even within familiar context, place fields in the AD model rats exhibit a significant decline in stability across days, impeding the formation of precise spatial maps through learning (Broussard, Redell et al. 2022). However, learning and memory can remain partially preserved despite these impairments in hippocampal spatial representations, a phenomenon consistent with cognitive resilience (Azarpazhooh, Avan et al. 2020, Ossenkoppele, Lyoo et al. 2020, Gómez-Isla and Frosch 2022, Chen, Pineau et al. 2025, Jia, Xu et al. 2025, Serino, Stramba-Badiale et al. 2025). This mismatch between impaired spatial representations and retained memory performance suggests that place field-based spatial coding may not explain retained memory performance in AD. Recent work suggests that hippocampal population activity also contains internally organized temporal cofiring structure, which can remain stable even when place fields drift or remapping (Levy, Carrillo-Segura et al. 2023, Fenton 2024). This raises the possibility that implicit temporal structures may retain experience-dependent plasticity in AD and contribute to memory function.

The integrity of this internal temporal structure is directly reflected during neural ensemble reactivation in the offline memory consolidation phase (Langdon, Genkin et al. 2023, Castelli, Lopes-Dos-Santos et al. 2025). Growing evidence indicates that such reactivation is not merely a simple replay of past behavioral trajectories but a spontaneous manifestation of intrinsic neural dynamic structures (Dragoi and Tonegawa 2011). Crucially, reactivation sequences are not constrained by specific behavioral histories but generate stochastic and continuous neural trajectories based on spatial connectivity (Stella et al., 2019). Furthermore, reactivation can flexibly generate novel detour paths based on contextual changes even when explicit place fields have not yet remapped (Widloski and Foster 2022). These findings suggest that reactivation is not merely a passive record of external experience, but also be shaped by intrinsic temporal structures. However, in AD, this intrinsic temporal organization may also be disrupted during subsequent offline memory consolidation. A key insight is derived from a previous study finding that neurons in AD mice maintain pathologically rigid firing sequences across distinct contexts, even when explicit place fields are impaired (Cheng and Ji, 2013). Based on this, we hypothesize that the AD brain may retain internal temporal structures.

In this study, we aimed to investigate whether a functional dissociation exists between deficits in explicit spatial representation and implicit cofiring dynamic in AD. To assess how these representations evolve during long-term experience, we monitored population activity in the hippocampal CA1 region of App^NL-G-F^ rats using a cross-day familiar-novel context alternating task. As rats gained experience across days, the App^NL-G-F^ group consistently displayed severe impairments in explicit remapping, with spatial coding failing to refine with experience. In contrast, implicit cofiring dynamics gradually strengthened with experience and developed the capacity to discriminate between distinct contexts. Further analysis revealed that this improvement of implicit coding was associated with an aberrant enhancement of reactivation consistency during sharp wave-ripples (SWRs) in rest periods. These findings reveal an explicit–implicit dissociation in the AD hippocampal network, suggesting that experience-dependent refinement of implicit temporal coding may support learning-related hippocampal network dynamics despite impaired spatial remapping. This framework may further shed light on neuromodulation strategies addressing the re-establishing the precise registration of internal temporal dynamics to external cues in AD.

## Results

### Impaired contextual discrimination during repeated exposure in App^NL-G-F^ rats

To investigate neural mechanisms of integrating novel sensory inputs into intrinsic neural representations and making discrimination in Alzheimer’s disease, we recorded hippocampal CA1 activity in 8-10 month-old App^NL-G-F^ rat. This model exhibits significant Aβ plaques and hyperphosphorylated tau protein, but does not yet show widespread neurodegeneration, characteristic of the early-to-mid stages of AD (Pang, Jiang et al. 2022). We utilized a familiar-novel context alternating task, which consisted of four 20-minute exploration sessions of either a familiar (A1/ A2) or a novel (B1/ B2) context, separated by 10-minute rest intervals (Fig. 1A). We first observed that App^NL-G-F^ rats exhibited weakened global remapping capability compared to WT rats (Fig. 1B, C; linear mixed effects model, spatial correlation, t(468) = 2.3, p = 0.02, β = 0.2; population vector correlation (PVC), t(8616) = 2.6, p = 0.009, β = 0.3; rate overlap, t(471) = 5.3, p = 2.0×10^-7^, β = 0.2). However, by considering the first and second A-B transitions (i.e., A1-to-B1 and B2-to-A2) separately, we found that context discriminative ability was reduced in App^NL-G-F^ rats during the second transition, indicated by an increase of across-session representational similarity (Fig. 1B, D; App^NL-G-F^ vs. 0, linear mixed effects model, spatial correlation, t(145) = 5.0, p = 1.9×10^-6^, β = 0.1; PVC, t(1799) = 2.0, p = 0.049, β = 0.1; rate overlap, t(135) = 3.5, p = 7.4×10^-4^, β = 0.1), with App^NL-G-F^ rats exhibiting a significantly greater increase than WT (WT vs. App^NL-G-F^, spatial correlation, t(233) = 2.1, p = 0.04, β = 0.07; PVC, t(3701) = 2.2, p = 0.03, β = 0.2; rate overlap, t(169) = 2.8, p = 0.005, β = 0.1). In contrast, A1A2 spatial similarity was comparable between groups, indicating preserved within-context stability and suggesting that the B2A2 deficit was not driven by general within-day drift (Figure 1–figure supplement 3). These findings suggest that contextual discrimination deficits in App^NL-G-F^ rats are dynamic and state-dependent over time. To test whether the reduced remapping ability during the B2-to-A2 transition was linked to a specific ensemble, we detected place cells that showed stable spatial firing during the within-context A1A2 comparison (spatial correlation above the 60th percentile of the A1A2 distribution) but exhibited insufficient remapping during the B2A2 transition (spatial correlation above the 95th percentile of the A1B1 distribution), and defined them as “confused cells”. This criterion allowed us to quantify changes in environmental discriminative ability across the two sequential transitions. We found that the proportion of confused cells reached 18.9% in App^NL-G-F^ rats, which was much higher than 3.4% in WT rats (Fig. 1E, Chi-square test, χ^2^(1,179) = 10.9, p = 9.8×10^-4^).

**Figure 1.**
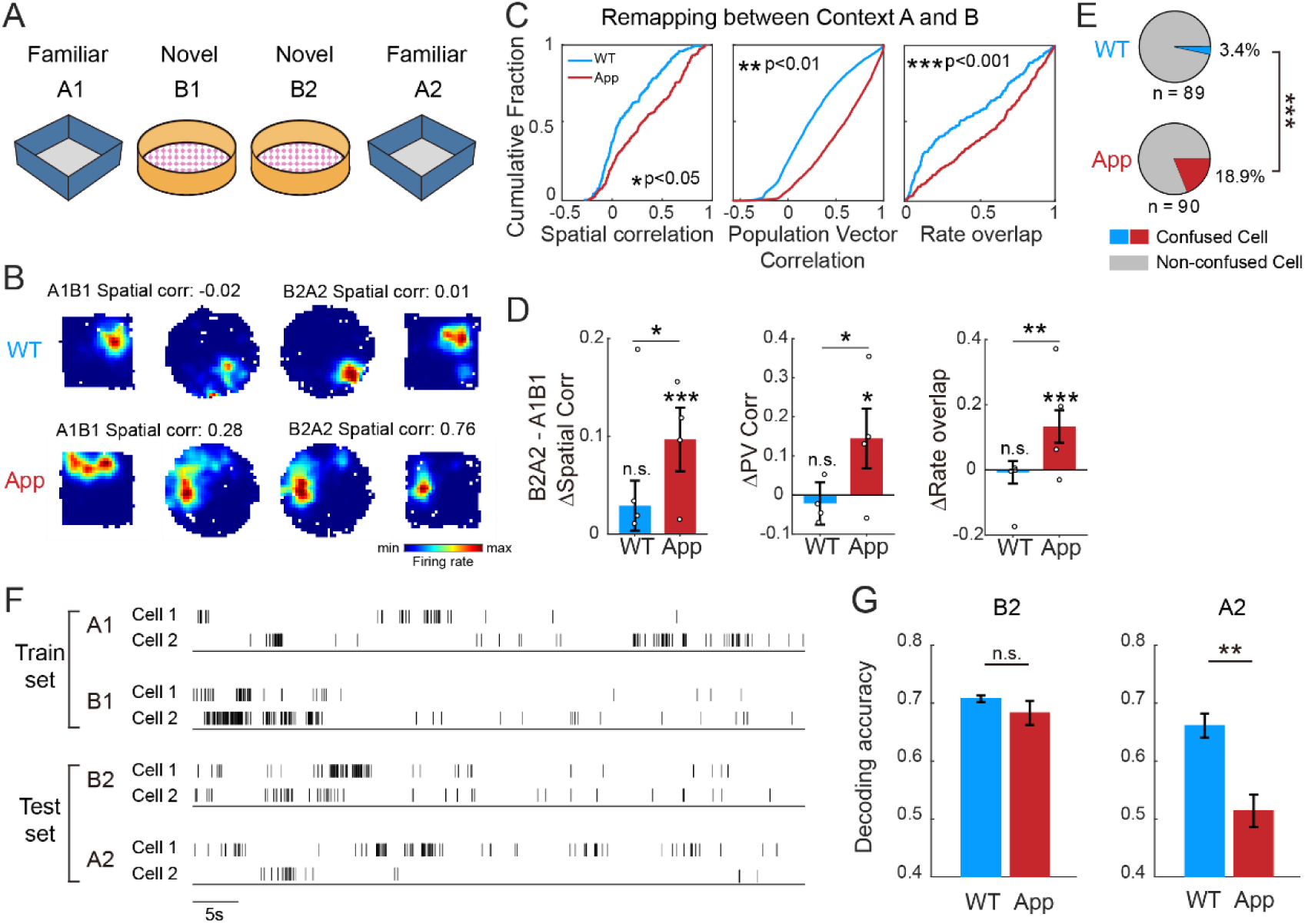
Single-cell spatial confusion upon returning to familiar context in App^NL-G-F^ rats. **A.** Schematic of the familiar-novel context alternating task. **B.** Representative rate maps of CA1 place cells across sessions. **C.** Quantification of remapping during the transition between context A and context B. Left: Spatial correlation. Middle: Population vector correlation. Right: Rate overlap. **D.** Difference in the degree of remapping between the initial transition from context A to context B (A1–B1) and the return transition from context B to context A (B2–A2). Left: ΔSpatial correlation. Middle: ΔPopulation vector correlation. Right: ΔRate overlap. **E.** Proportion of confused cells in the total population. Confused cells were defined as neurons with A1A2 spatial correlations above the 60th percentile of the A1A2 distribution and B2A2 spatial correlations above the 95th percentile of the A1B1 distribution. **F.** Representative raster plots of place cell spike trains. **G.** Left: SVM decoding accuracy within session B2. Right: SVM decoding accuracy within session A2. Data are presented as mean ± SEM.

Given that neural manifolds derived from neuronal spike trains can represent different contexts or task strategies (Tang, Shin et al. 2023), we further hypothesized that the temporal structure of spike trains contained distinct contextual information. To test this hypothesis, we employed a support vector machine (SVM) decoder as a quantitative readout of contextual discriminability, training it on spike train activities from A1 and B1 and testing its performance on B2 and A2 across repeated context exposures (Fig. 1F). We found that the decoding accuracy for B2 remained high in both groups (Fig. 1G, Student’s t-test, t (163) = 0.72, p = 0.48, Cohen’s d = 0.1), however, the decoding accuracy for A2 was significantly reduced in App^NL-G-F^ rats compared to WT rats (Fig. 1G, Student’s t-test, t (163) = 3.1, p = 0.002, Cohen’s d = 0.7). These results suggest that neural ensembles in App^NL-G-F^ rats could respond to the novel information during the initial exposure, while they lost the ability of instantaneous shifting during repeated context transitions.

This raised the question of whether the retrieval deficit arose from unstable spatial representations or from a disruption in the intrinsic temporal organization of the neural circuit. To address this issue, we introduced a theoretical framework showing that remapping is essentially the projection of an invariant, high-dimensional neural manifold onto different low-dimensional environments (Fenton 2024). The remapping analyses mainly referred to place coding based on temporal-independent activity that the firing rate of single cells directly represents the physical location of the animal in the external context (Fig. 2A). In contrast, the temporal cofiring relationship between a pair of cells reflects the intrinsic sequential organization within the network, which could be measured by Kendall’s τ (Neymotin, Talbot et al. 2017, Levy, Carrillo-Segura et al. 2023). The pairwise Kendall’s τ was calculated for all cell pairs, thereby we assessed the correlation of these τ vectors across sessions (Neymotin, Talbot et al. 2017) and obtained a metric termed population cofiring correlation (PCC) (Fig. 2A). During the initial A1-B1 transition, App^NL-G-F^ rats maintained a low PCC comparable to WT rats, with low similarity of cofiring patterns between familiar and novel contexts (Fig. 2B, C, linear mixed effects model, linear mixed effects model, A1B1, F(1, 4) = 0.03, p = 0.9, β = 0.02, n_WT_ = 3 rats, n_App_ = 3 rats). The distinct contexts are encoded through the reorganization of these temporal patterns, where certain cell pairs switch to opposing (anti-cofiring) modes to discriminate different contexts. However, during the second B2-A2 transition, temporal representation of App^NL-G-F^ rats exhibited significantly higher correlation between different contexts compared to that of WT rats (Fig. 2B, C, F(1, 4) = 9.5, p = 0.04, β = 0.2). In contrast, PCC between same contexts remained stable in App^NL-G-F^ rats (Fig. 2C, F(1, 4) = 1.1, p = 0.3, β = 0.09). These results were consistent with the global remapping findings, suggesting that the neural representation of App^NL-G-F^ rats lost the capacity to distinguish the familiar context from the recent novel context during repeated exposure.

**Figure 2.**
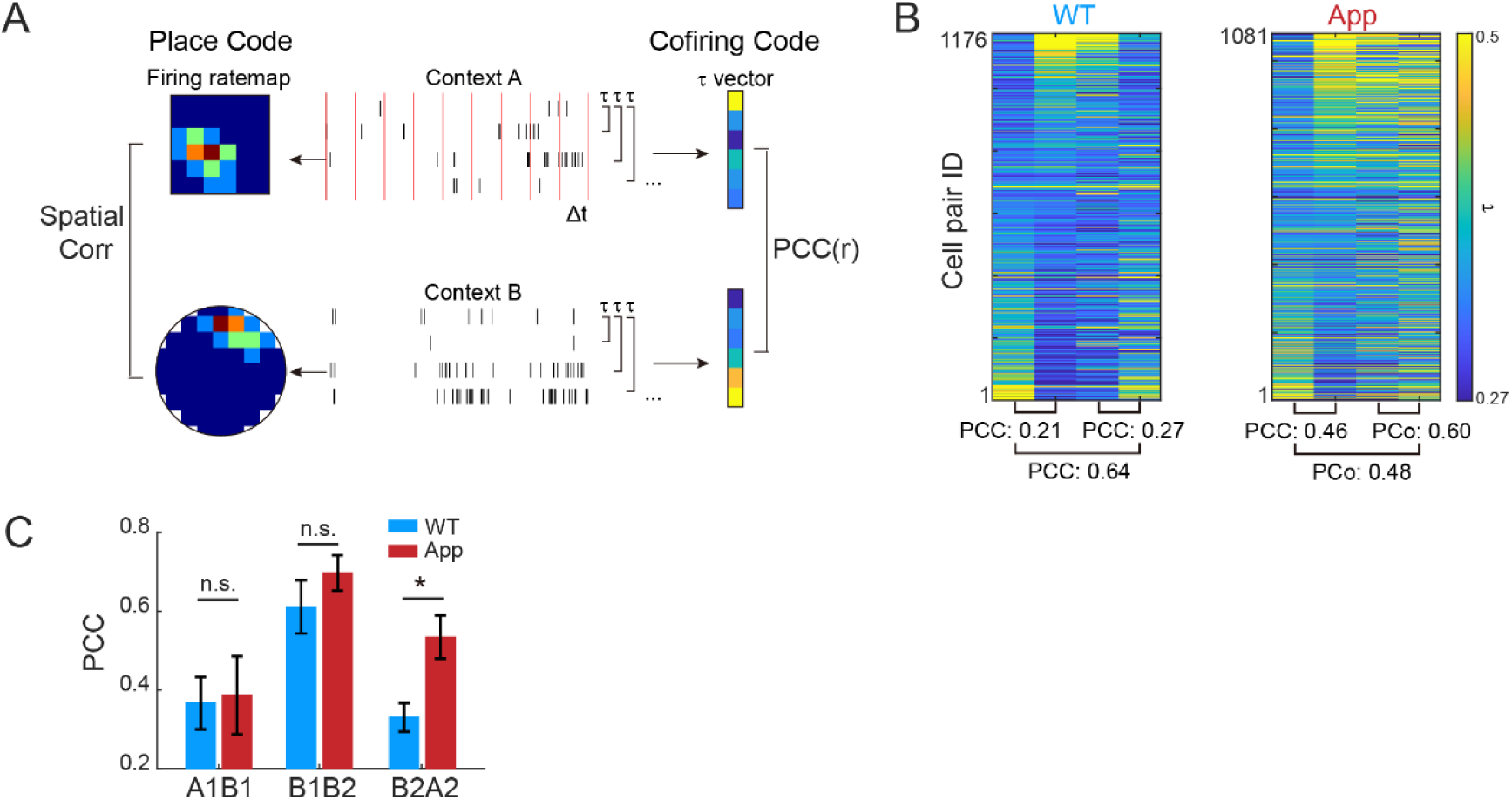
Implicit pairwise cofiring fails to discriminate between contexts B2 and A2 in App^NL-G-F^ rats during the first experimental day. **A.** Schematic of explicit place code and implicit cofiring code. Left: A single place cell forms a spatial rate map, discarding temporal information. Right: Kendall’s rank correlation (τ) is calculated between spike trains of cell pairs, incorporating temporal information. **B.** Representative heatmaps of pairwise τ values in each session. The bottom number shows the correlation of τ vectors across sessions, defined as the Population Cofiring Correlation (PCC). **C.** PCC between different sessions.

### Cofiring coding within ensemble supports experience-dependent contextual discrimination in App^NL-G-F^ rats

To investigate how experience-dependent neural coding changes in AD rats during repeated exposure to distinct contexts, we performed the novel context transition task for six consecutive days and measured cofiring dynamics within ensemble over time. The PCC between sessions in the same context (i.e., A1A2, B1B2) and in the different contexts (i.e., A1B1, B2A2) were obtained over 6 days in both genotypes (Fig. 3A). We found that the temporal correlation within the same context was significantly higher than that across different contexts in App^NL-G-F^ rats, as in WT rats (Fig. 3B, two-way ANOVA, main effect of context type, WT F(1, 48) = 316.3, p = 9.1×10^-23^, partial η^2^ = 0.9, App^NL-G-F^ F(1, 60) = 24.9, p = 5.5×10^-6^, partial η^2^ = 0.3). Importantly, the PCC within the same context and across different contexts exhibited divergent trends over days, indicating an experience-dependent development of contextual recognition in both groups (Fig. 3B). We then defined ΔPCC as the difference in PCC between the same and different contexts, and found that ΔPCC in App^NL-G-F^ rats increased across days similarly to that in WT rats (Fig. 3A, C, linear regression, WT, r = 0.4, p = 0.02, App^NL-G-F^, r = 0.3, p = 0.04), but was lower than that in WT rats (two-way ANOVA, main effect of group, F(1, 54) = 15.4, p = 2.5×10^-4^, partial η^2^ = 0.2).

**Figure 3.**
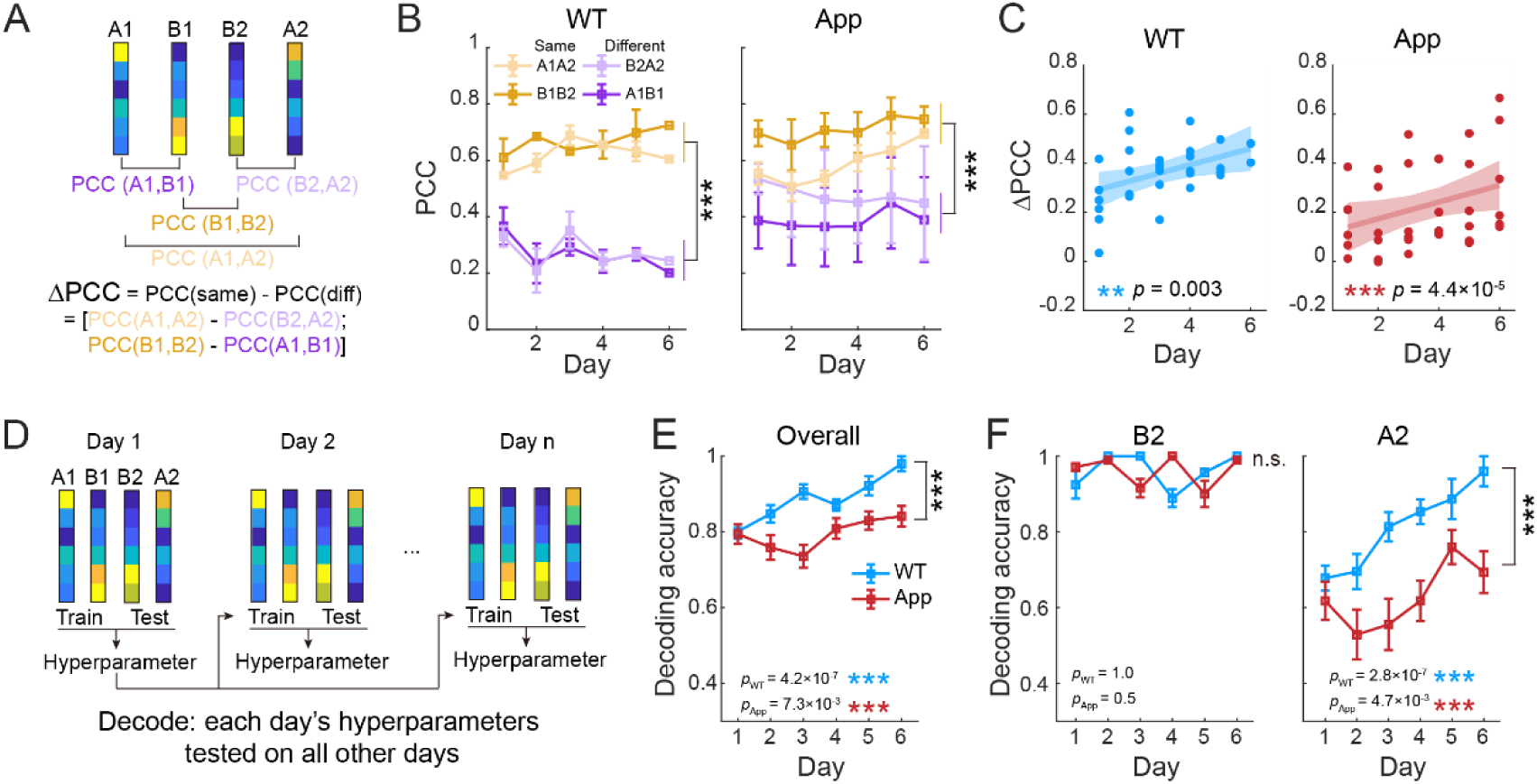
Cross-day pairwise cofiring gradually discriminates distinct contexts in App^NL-G-F^ rats. **A.** Schematic of PCC analysis across sessions. Purple: Cross-context correlations (Dark purple: A1B1; Light purple: B2A2). Yellow: Same-context correlations (Dark yellow: B1B2; Light yellow: A1A2). The discrimination index (ΔPCC) is defined as the difference between same-context and cross-context PCCs. **B.** Evolution of PCC values across training days. **C.** Evolution of ΔPCC across training days. **D.** Schematic of the SVM decoding workflow. τ vectors from A1 and B1 served as the training set, while those from B2 and A2 served as the testing set. To ensure unbiased comparison, optimal hyperparameters were identified for each day and cross-validated across all days. **E.** Overall SVM decoding accuracy for combined B2 and A2 sessions. **F.** Left: Accuracy specific to session B2. Right: Accuracy specific to session A2.

To further test whether cofiring structure varying over time supports context recognition during repeated exposures (i.e., B2 and A2), we trained an SVM classifier using τ vectors from A1 and B1 as the training set, and evaluated decoding performance for B2 and A2 across days (Fig. 3D). We found that decoding based on the cofiring structure effectively discriminated between different contexts, achieving a decoding accuracy of 98% in WT and 84% in App^NL-G-F^ rats. Although the decoding accuracy was significantly lower in App^NL-G-F^ rats (Fig. 3E; two-way ANOVA, F(1,483) = 37.1, p = 2.2×10^-9^, partial η^2^ = 0.07, n_WT_ = 225, n_App_ = 270 cross-validation tests), both WT and App^NL-G-F^ showed experience-dependent improvement (linear mixed effects model, WT, F(1, 223) = 27.2, p = 4.2×10^-7^, β = 0.03; App^NL-G-F^, F(1, 268) = 7.3, p = 7.3×10^-3^, β = 0.02). Interestingly, by separately decoding two different contexts, we found that decoding accuracy for B2 remained high (>90%) across days for both genotypes and showed no significant change with experience (Fig. 3F, left, two-way ANOVA, F(1,483) = 0.006, p = 0.9, partial η^2^ = 1.2×10^-5^; linear mixed effects model, WT, F(1, 223) = 0.002, p = 1.0, β = -2.9×10^-4^; App^NL-G-F^, F(1, 268) = 0.4, p = 0.5, β = -0.003, n_WT_ = 225, n_App_ = 270 cross-validation tests). In contrast, the recognition for A2 improved gradually over time with experience in both groups (Fig. 3F, right; linear mixed effects model, WT, F(1, 223) = 28.1, p = 2.8×10^-7^, β = 0.06; App^NL-G-F^, F(1, 268) = 8.1, p = 4.7×10^-3^, β = 0.03, n_WT_ = 225, n_App_ = 270 cross-validation tests), reaching a maximum of 96% in WT rats and 76% in App^NL-G-F^ rats (two-way ANOVA, F(1,483) = 36.5, p = 5.0×10^-9^, partial η^2^ = 0.07).

One possible explanation for this finding is that the spatial representations of hippocampal place cells develop over time, becoming more distinct across contexts as experience accumulates (Lever, Wills et al. 2002, Hainmueller and Bartos 2018, Plitt and Giocomo 2021). To test this hypothesis, we then measured the place field size of each cell as a low-dimensional place coding. Over time, place fields became significantly smaller in WT rats but not in the App^NL-G-F^ group (Figure 3–figure supplement 1, two-way ANOVA, group × day interaction, F(1,989) = 6.4, p = 0.01, partial η^2^ = 0.006, post hoc comparison, WT, p = 5.0×10^-5^, App^NL-G-F^, p = 0.3). These findings indicate that although place coding fails to update with experience accumulates, the internal cofiring structure at temporal scale exhibited increasing specificity across contexts, which may support the emerging of context discrimination in App^NL-G-F^ rats.

### Environment discrimination requires implicit temporal coding rather than explicit spatial coding in App^NL-G-F^ rats

The measurement of cofiring coding within ensembles involves information from both explicit spatial coding and implicit temporal coding. By detecting cell pairs with overlapping place fields, we observed that greater place field overlap was associated with stronger cofiring, and vice versa (Fig. 4A). We found that the τ difference between contexts A and B was positively correlated with the place field overlapping difference between them (Fig. 4B, linear regression, WT, r = 0.7, p = 2.1×10^-30^, App^NL-G-F^, r = 0.6, p = 5.9×10^-38^). However, the proportion of variance (R^2^) in cofiring structure explained by spatial overlap was significantly lower in App^NL-G-F^ rats than in WT rats (Fig. 4C, linear mixed effects model, F(1, 32) = 10.7, p = 0.003, β = -0.1, n_WT_ = 15 sessions, n_App_ = 19 sessions). These results indicate that the relationship between spatial overlap and temporal cofiring remained detectable in App^NL-G-F^ rats, but was significantly weaker than in WT rats. To further determine which would be essential for the cofiring coding development with experience, we separately measured the effect of these two factors on the environment discrimination. We found that spatial correlation between different environments was significantly lower in App^NL-G-F^ rats than that in WT rats, which difference was persistent across days (Fig. 4D-E, linear mixed effects model, main effect of group, F(1, 2674) = 4.0, p = 0.046, β = 0.2, n_WT_ = 1006 cells, n_App_ = 1682 cells), as well as the population vector correlation between different environments (Fig. 4F, linear mixed effects model, main effect of group, F(1, 22) = 5.5, p = 0.03, β = 0.3, n_WT_ = 15 sessions, n_App_ = 19 sessions). When A1-B1 and A2-B2 epoch pairs were analyzed separately, neither spatial correlation nor PVC showed a significant change across days (Figure 4–figure supplement 1).

**Figure 4.**
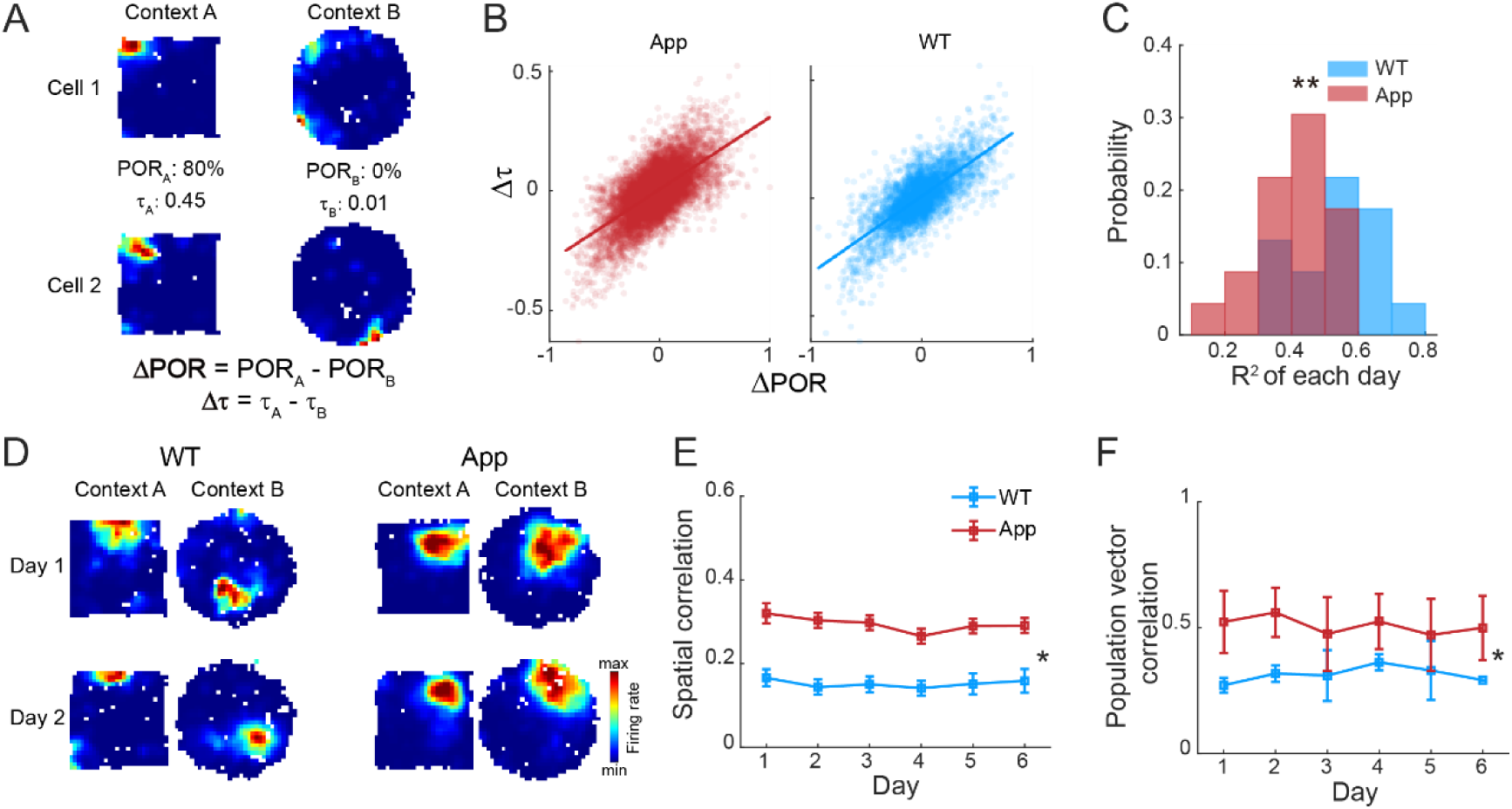
Explicit spatial discrimination stagnates and decouples from cofiring dynamics in App^NL-G-F^ rats. **A.** Representative rate maps of a cell pair. Left: Rate maps in context A. POR_A_ denotes the place field overlap ratio, and τ_A_ denotes the pairwise Kendall’s tau in context A. Right: Same for context B. ΔPOR and Δτ denote the differences between contexts. **B.** Scatter plot of ΔPOR versus Δτ for all cell pairs. R^2^ indicates the goodness of fit, reflecting the predictive power of spatial overlap for temporal cofiring. **C.** Distribution of daily goodness of fit (R^2^). **D.** Representative rate maps of place cells across training days for WT and App^NL-G-F^ rats. **E.** Spatial correlation between context A and B across training days. **F.** Population vector correlation between context A and B across training days.

To further dissociate cofiring dynamics from spatial tuning, i.e. the implicit temporal coding, we computed the position-tuning independent rate (PIR) for each neuron. PIR was obtained by subtracting the expected rate from the observed rate, where the expected rate was based on the cell’s spatial rate map at the animal’s current position (Fig. 5A). Because PIR depends on expected rates derived from spatial rate maps, we also assessed how well spatial tuning explained neuronal firing using the spatial explained variance (SEV). SEV did not differ significantly between genotypes, indicating that spatial tuning explained a comparable fraction of firing-rate variance in WT and App^NL-G-F^ rats (Figure 5–figure supplement 2). We next calculated τ based on PIR activity (Fig. 5B), and found that the implicit temporal coding during repeated exposure to the familiar environment (A2) was similar to that during B2, as reflected by a trend toward higher PCC for B2-A2 in App^NL-G-F^ rats (Fig. 5C, linear mixed effects model, F(1, 32) = 4.0, p = 0.053, β = 0.1, n_WT_ = 15 sessions, n_App_ = 19 sessions), consistent with the cofiring pattern observed in Fig. 2. We then trained an SVM decoder by PIR-based τ vectors and performed decoding between environments A and B. We found that the decoding accuracy based on implicit temporal coding in App^NL-G-F^ rats was significantly increased with experience as that in WT rats (Fig. 5D, linear mixed effects model, WT, F(1, 223) = 4.2, p = 0.04, App^NL-G-F^, F(1, 268) = 18.4, p = 2.3×10^-5^), although lower in App^NL-G-F^ rats than in WT rats (linear mixed effects model, main effect of group, F(1, 491) = 7.6, p = 0.006, β = -0.2, n_WT_ = 225, n_App_ = 270 cross-validation tests). By contrast, decoding based on τ calculated from expected firing rates showed no significant experience-dependent change in App^NL-G-F^ rats (Figure 5–figure supplement 1). These findings suggest that the implicit temporal coding supports the environment discrimination developed with experience in App^NL-G-F^ rats, which was independent of explicit spatial coding impairment.

**Figure 5.**
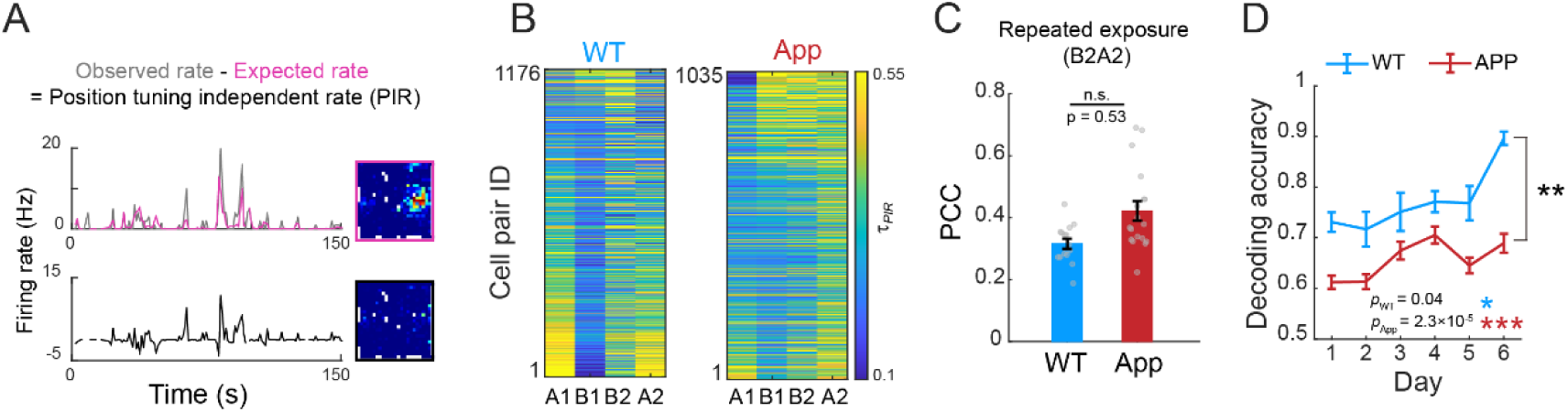
Position-independent cofiring dynamics evolve to discriminate contexts with experience. **A.** Schematic of the Position Tuning Independent Rate (PIR) calculation. The expected rate (pink line) is derived by projecting the animal’s real-time position onto the spatial rate map (pink box), which displays clear place fields. The observed rate (gray line) is the actual firing rate. All rate vectors were computed in 100 ms bins. PIR is defined as the subtraction of the expected rate from the observed rate. Re-mapping the PIR onto spatial coordinates (black box) confirms the absence of place fields. **B.** Representative heatmaps of pairwise τ*_PIR_* calculated using PIR. **C.** PCC of τ*_PIR_* during repeated exposure (i.e., B2A2). **D.** SVM decoding accuracy using τ*_PIR_* vectors. The decoding strategy follows the workflow outlined in Figure 3D.

### Enhanced reactivation consistency during offline consolidation in App^NL-G-F^ rats

The temporal organization of hippocampal firing within theta cycles during active exploration and within SWRs during resting was considered essential for constructing coherent spatial representations (Tang and Jadhav 2022, Ji, Chu et al. 2026). To first investigate whether the preserved implicit temporal coding relies on theta rhythmic coordination, we analyzed the time lag between spike trains of cell pairs (Fig. 6A). We calculated the cross-correlograms (CCG) in both directions to avoid temporal ambiguity caused by place field traversals on different directions (Fig. 6B). Theta index was measured to quantify to what extent cofiring patterns were modulated by theta rhythm (Fig. 6C). We found that the place cells in both WT and App^NL-G-F^ rats showed comparable theta modulation (Fig. 6D, linear mixed effects model, F(1, 36293) = 1.4, p = 0.2, β = 3.3×10^-3^, n_WT_ = 7328 cell pairs, n_App_ = 28967 cell pairs), indicating that the dissociation between explicit and implicit coding is not likely driven by theta entrainment in App^NL-G-F^ rats.

**Figure 6.**
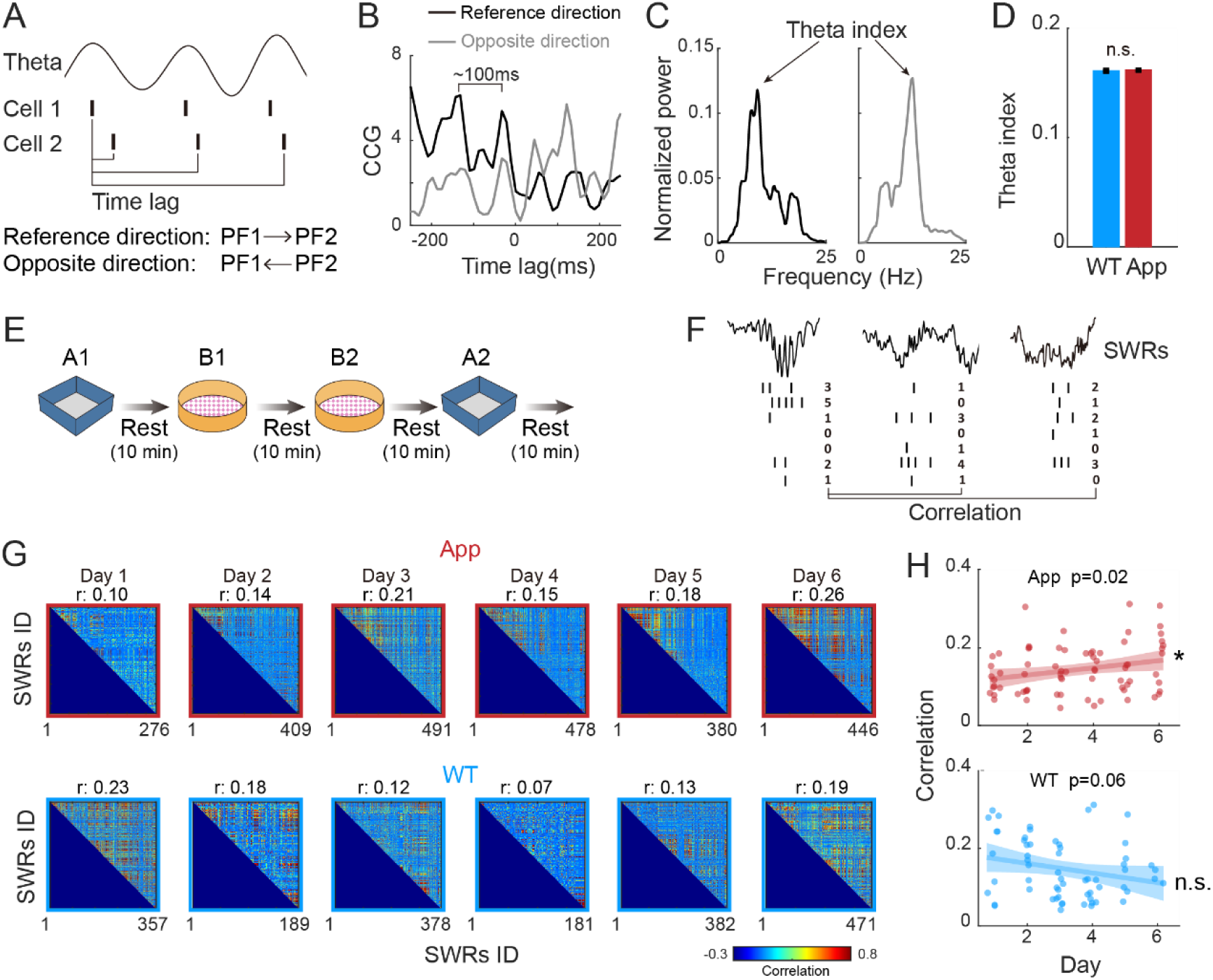
Progressive increase in reactivation consistency during rest periods in App^NL-G-F^ rats. **A.** Schematic illustration showing the coordination of cell pair firing by theta rhythms. The time lag between spikes is analyzed across two running directions: from place field 1 (PF1) to PF2 (Reference direction) and from PF2 to PF1 (Opposite direction). **B.** Representative cross-correlograms (CCG) of a cell pair are plotted as a function of the time lag between spikes. The black and gray lines represent movement in the reference and opposite directions, respectively. Note that the periodicity of the peaks resembles the theta rhythm cycle (∼100 ms). **C.** Frequency power spectra of the CCG shown in B. The peak value within the theta band (6-12 Hz) is defined as the theta index. **D.** Quantification of theta indices across all recorded cell pairs. **E.** Schematic of the experimental timeline showing the behavior task in which rats rested for ten minutes between each running session. **F.** Schematic illustrating the computation of neuronal reactivation consistency. For each SWR event, a population vector was constructed consisting of the spike counts from all recorded neurons. Reactivation consistency was then quantified by computing the correlation between these population vectors across different SWR events. **G.** Representative examples showing the changes in reactivation consistency across training days for a WT rat and an App ^NL-G-F^ rat. **H.** Group data quantifying the changes in reactivation consistency across training days.

We next focused on offline resting state which offered a unique opportunity to assess the brain’s intrinsic generative models, as neural reactivation is driven by internal network dynamics in the absence of external sensory cues (Widloski and Foster 2022, Zaki, Pennington et al. 2025). We detected SWRs and quantified the consistency among SWR-associated reactivation patterns, with higher event-to-event correlations suggesting more stable attractor-like dynamics, consistent with the network repeatedly returning to similar ensemble states (Fig. 6F). We found that the experience-dependent change in reactivation consistency differed between App^NL-G-F^ and WT rats, (Fig. 6G, H; linear mixed effects model, group × day interaction, F(1, 129) = 8.6, p = 0.004, β = 0.02, n_WT_ = 61 sessions, n_App_ = 72 sessions). Reactivation consistency increased significantly across days in App^NL-G-F^ rats (linear mixed effects model, F(1, 70) = 6.1, p = 0.02), whereas no significant increase was observed in WT rats (F(1, 59) = 3.7, p = 0.06). These changes in reactivation consistency were not accompanied by differences between genotypes in SWR incidence, SWR duration, participating cells per event, or mean rest firing rate (Figure 6–figure supplement 1). It suggests that App^NL-G-F^ rats progressively strengthen their intrinsic network structure across days despite the fragility of their explicit spatial maps. The enhanced reactivation consistency aligns with the improvement in implicit dynamics observed during behavior, suggesting a potential compensatory role for internally organized network activity in App^NL-G-F^ rats.

## Discussion

In the present study, by examining the evolution of hippocampal neural ensemble activity across multiple days, we found a dissociation between preserved implicit temporal structures and impaired explicit spatial representation during novel context experience in early-stage AD pathology. Although place field remapping was disrupted, implicit cofiring dynamics remained capable of developing with experience. Moreover, SWR-associated reactivation during offline rest showed enhanced pattern consistency, suggesting that offline network activity may help stabilize these internal temporal structures in AD. Together, these findings suggest that AD selectively impairs the registration of hippocampal activity to external spatial cues, while preserving the implicit temporal structures that may support the experience-dependent acquisition of contextual discrimination.

We found that App^NL-G-F^ rats exhibited impaired explicit spatial representations during context transitions, reflected in reduced place cell remapping across environments, consistent with previous single neuron studies (Jun, Bramian et al. 2020, Broussard, Redell et al. 2022). We further revealed that this deficit in spatial discrimination was persistent and failed to improve despite the accumulation of long-term learning experience. However, single-cell coding deficits alone may not fully account for the residual function of hippocampal networks in AD. The recent reregistration theory proposed by Fenton offers a critical perspective on this issue (Fenton 2024). This theory posits that the core of hippocampal neural coding lies not in passive responses to the external context, but in maintaining a neurocentric internal cofiring pattern. Typically, temporal cofiring relationships among large populations of cells are highly robust and operate independently of external contextual changes, while distinct contexts are differentiated primarily by the activity of anti-cofiring subpopulation (Levy, Carrillo-Segura et al. 2023). Within this framework, we found that although App^NL-G-F^ rats failed in explicit remapping, their implicit cofiring dynamics retained the capacity to demonstrate increased discrimination between distinct contexts with accumulated experience. This demonstrates that the intrinsic organization of the AD network remains preserved. Consequently, cognitive deficits stem from a pathological dissociation, wherein this intact internal structure fails to correctly register to external spatial cues.

The preservation of such intrinsic network organization is further substantiated by evidence from theta oscillations. We found that the capacity of theta rhythms to coordinate pairwise neuronal firing remains intact, implying the persistence of a fundamental temporal framework driven by internal mechanisms. This phenomenon can be interpreted through the multiplexed phase coding hypothesis (Sueoka et al., 2025), which suggests that hippocampal theta rhythms integrate two distinct information streams: spatial coding dependent on external cues, and temporal structure derived from internal dynamics. Notably, the component driven by internal cues is characterized by high robustness, whereas the component dependent on external cues appears relatively fragile. This contrast between robust internal dynamics and fragile external coding is corroborated by findings in the rTg4510 Tau pathology model, where hippocampal ensembles were observed to revert to a state dominated by internal dynamics when external spatial cues failed to effectively drive the network, manifesting as rigid firing sequences maintained across distinct contexts (Cheng and Ji 2013). Taken together, these prior findings and our results suggest that the AD brain may lose the ability to register representations to external spatial coordinates, while partially retaining intrinsic temporal structures driven by network dynamics.

We found that, as experience accumulated across days, App^NL-G-F^ rats exhibited progressively increased reactivation consistency during SWRs. Notably, this change paralleled the experience-dependent increase in contextual discrimination by implicit temporal structure during active exploration, while explicit spatial representations remained impaired. This change may reflect a compensatory reorganization of cellular reactivation during awake rest. One possible explanation is that AD-associated hyperexcitability during low-arousal states, such as sleep or rest, may influence reactivation dynamics (Zarhin, Atsmon et al. 2022). Given the impairment of explicit spatial representations, altered reactivation patterns may reflect reduced flexibility in updating external contextual representations, potentially leading to more stereotyped internal manifolds (Fenton 2024). Therefore, the increased discrimination capacity of implicit cofiring dynamics observed in App^NL-G-F^ rats may represent a potential compensatory stabilization of internal network dynamics during SWRs despite impaired spatial representations.

Several limitations of the present study remain to be addressed in future work. First, while impaired explicit state transitions likely involve compromised network resetting mediated by PV+ interneurons (Hijazi et al., 2020), our inability to distinguish inhibitory subtypes prevented a direct mechanistic analysis at the microcircuit level. Second, the interpretation of the SWR reactivation results remains limited. The limited number of recorded units prevented us from determining whether increased reactivation similarity reflected enhanced memory consolidation or reduced ensemble diversity. Moreover, the causal relationship between SWR reactivation and cofiring structures requires further verification. Future studies should employ closed-loop optogenetics to disrupt SWRs, thereby testing whether this compensatory reactivation is essential for maintaining implicit network function. Additionally, how downstream regions like the prefrontal cortex interpret these dissociated codes, where intact implicit signals coexist with corrupted spatial maps, remains a critical open question. Whether these findings generalize across sexes remains to be determined, as this study was performed exclusively using male rats.

Finally, our findings offer a novel perspective for clinical intervention. Rather than focusing solely on rescuing damaged explicit memory, future neuromodulation strategies might target the “re-registration” of preserved internal dynamics to external cues. By utilizing these robust intrinsic structures as anchors, such interventions could open a new therapeutic window for cognitive rehabilitation in AD.

## Acknowledgments

This work was supported by grants from the National Key Research and Development Program of China (2024YFF1206500 to C.Z.), the National Natural Science Foundation of China (T2322021, 82271218 to C.Z., 82501763 to Y.W.), and the Natural Science Foundation of Tianjin (25JCJQJC00290 to C.Z.).

## Author contributions

C.Z., X.W., and K.P and designed the experiment; X.W. and Y.W. carried out the electrophysiological recordings and behavioral task; X.W. and C.Z. designed analyses; X.W. wrote analysis programs and analyzed the data; X.W. and C.Z. wrote the paper; and C.Z. supervised the project. All authors discussed the results.

## Materials and methods

### Subjects

The School of Pharmaceutical Sciences at Tsinghua University kindly donated animals for this study. 8-10 months old male App^NL-G-F^ rats (n=4) and WT rats (n=4) were used in this research. Rats were maintained in standard housing conditions before surgery. After surgery, rats were housed in custom-built acrylic cages (40cm*40cm*40cm) on a reverse light cycle (Light: 9pm to 9am; Dark: 9am to 9pm). Active waking behavior recordings were performed during the dark phase of the cycle. Rats were pre-trained to perform the open field exploration before recording drive implantation surgery. All experiments complied with the Animal Care and Use Committee guidelines of the Tianjin Hospital of Tianjin University.

### Recording drive implantation and surgery

Each drive contained 21 independently moveable tetrodes fabricated from 17 μm polyimide-coated platinum-iridium wires (plated to ∼150–300 kΩ). Drives were implanted above the bilateral hippocampus (AP -3.8 mm, ML ±2.5 mm) and anchored with dental cement and skull screws. Eighteen tetrodes targeted the CA1 stratum pyramidale, one targeted the apical dendritic layer, and one served as a cortical reference. The number of cells recorded per day is summarized in Table S1 and Figure 1–figure supplement 2A.

### Histological verification of tetrode locations

Following completion of electrophysiological recordings, animals were deeply anesthetized with urethane and transcardially perfused with phosphate buffered saline (PBS), followed by 4% paraformaldehyde. Brains were removed and post-fixed in 4% paraformaldehyde overnight at 4°C, followed by cryoprotection in a sucrose gradient (15% and 30%). Coronal brain sections (30 μm thickness) were prepared using a cryostat and stained with cresyl violet to visualize hippocampal structures and tetrode tracks. Images were acquired using a bright-field microscope, and tetrode tracks were examined to verify recording locations within the CA1 pyramidal cell layer. Representative sections are shown in Figure 1–figure supplement 1.

### Behavioral task

After recovering from surgery, rats performed a behavioral task in two distinct contexts constructed from reversible steel plates. The familiar context (Arena A) was a 100cm × 100cm cm blue square on gray vinyl flooring, fully explored pre-surgery (>4 sessions). The novel context (Arena B) was a 12-sided yellow enclosure (approximating a circle, 400 cm perimeter) on pink/white flooring. The task followed an A-B-B-A sequence. Each 20-min session was interleaved with 10-min rest periods in a resting box. During the exploration sessions, rats moved freely for the first 5 minutes; for the remaining 15 minutes, popcorn was scattered to encourage continuous exploration. Between sessions, the arena was cleaned with 75% ethanol to eliminate olfactory cues. Running speed during open field exploration is shown in Fig. S8B.

### Data acquisition

Neural signals were acquired using a Digital Lynx system and Cheetah software (Neuralynx, Bozeman, MT). LFPs were sampled at 2 kHz and filtered between 0.1 and 500 Hz. For unit activity, signals were bandpass filtered (600–6,000 Hz), thresholded at 55 μV, and sampled at 32 kHz. Animal position was tracked at 25 fps via a red LED on the HS-27 headstage.

### Spike sorting

Spike events detected during the running session were manually classified using the graphical clustering interface of MClust 4.4 (A.D. Redish, University of Minnesota, Minneapolis). Spike clusters were sorted based on two-dimensional projections of three waveform features (peak amplitude, energy, and peak-to-valley temporal differences) across all four tetrode channels. Putative excitatory neurons were identified by meeting the following criteria: (1) waveforms exhibiting pyramidal cell-like morphology, (2) a minimum inter-spike interval of 1 ms (refractory period), and (3) clear separation of clusters from noise or neighboring clusters in at least three independent feature projections.

### Firing rate maps and place fields

Units with mean firing rates exceeding 15 Hz were classified as putative fast-spiking interneurons and excluded from analysis. To characterize firing fields, the position data were sorted into 4 cm × 4 cm bins. For the open field map, the path was smoothed with a 21-sample boxcar window filter (400 ms; 10 samples on each side). Firing rate distributions were then determined by counting the number of spikes in each bin as well as the time spent per bin. A firing field was considered a contiguous area comprising of at least four bins where the firing rate was above 20% of the peak rate. Only units with peak firing rates over 1 Hz and main field size over 9 cm during the running session were defined as putative neurons with spatial representation, which were included for further analysis.

### Spatial correlation

Spatial correlation was calculated to quantify the similarity of spatial firing patterns across sessions (Colgin, Leutgeb et al. 2010). For each cell, the Pearson correlation coefficient was computed between firing rates at spatially matched bins of rate maps from different sessions. Spatial bins were excluded from analysis if they were unvisited in either session or had an occupancy time of less than 150 ms. Cells with a peak firing rate <1 Hz were also excluded.

### Population vector correlation (PVC)

Population Vector Correlation (PVC) was used to assess the similarity of ensemble firing patterns between sessions (Colgin, Leutgeb et al. 2010). Specifically, we organized the firing rate maps of all recorded cells into a 3-dimensional matrix, where two dimensions represented spatial position (4 × 4 cm bins) and the third dimension represented cell identity. For each spatial bin, the vector of firing rates across all cells was defined as the population vector. The PVC was then calculated as the Pearson correlation coefficient between the population vectors at corresponding locations in two different sessions.

### Kendall’s tau (τ) and Population Cofiring Correlation (PCC)

Given the sparsity and discreteness of spike data, we used Kendall’s rank correlation coefficient (Kendall’s τ) to quantify the temporal co-firing relationships between a pair of cells(Levy, Carrillo-Segura et al. 2023). Kendall’s τ assesses the consistency of rank ordering rather than specific magnitude, making it highly robust to non-normally distributed data and time series with numerous ties (e.g., abundant 0 and 1). Specifically, the 20-minute recording from each session was binned into 1-s intervals to extract spike trains, and pairwise Kendall’s τ coefficients were computed for all cell pairs within each session. The PCC was then defined as the Pearson correlation coefficient between the corresponding τ vectors of different sessions. To ensure data quality, cells with fewer than 10 spikes in every session were excluded from the analysis, so one rat from each group was not included.

### SVM decoding

To decode context from explicit firing patterns, spike trains were binned into 1-s intervals to generate firing rate vectors. The training set consisted of the rate vectors from sessions A1 and B1, while vectors from B2 and A2 constituted the testing set. For each recording day, we optimized the SVM hyperparameters (regularization parameter C and kernel scale σ). These daily-optimized parameters were then evaluated on datasets from other days to perform a cross-session parameter validation.

For the decoding of implicit temporal correlations, pairwise Kendall’s τ was computed in 2-minute non-overlapping windows, yielding a feature matrix of 10 samples per session. Matrices from A1 and B1 formed the training set, with B2 and A2 matrices serving as the testing set. Hyperparameters were optimized within each day, followed by the same cross-session parameter validation procedure described above.

### Position independent rate (PIR)

We computed the PIR to assess temporal firing fluctuations independent of spatial location. The expected rate was first generated by projecting the animal’s instantaneous position (sampled at 100 ms) onto the cell’s spatial rate map. Both this position-predicted signal and the observed spike train were then binned into 1-s intervals. The PIR was defined by subtracting the expected rate from the observed rate within each 1-s bin, thereby removing the component of firing driven purely by the animal’s spatial location.

### Spatial Explained Variance (SEV)

To quantified the fraction of firing rate variance explained by the spatial model, we defined the Spatial Explained Variance (SEV):

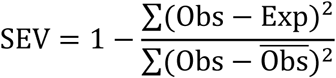

where Obs represents the observed firing rate and Exp represents the position-predicted expected rate. An SEV of 1 indicates that the observed firing rate is fully explained by the expected rate, whereas lower values indicate less variance explained by the spatial model.

### Direction-specific Cross-Correlograms (CCG)

To characterize the strength of temporal coordination between neuronal pairs by theta rhythms, we computed CCG based on the spike timing differences between a pair of cells. The CCG were constructed using a 5 ms bin size within a time lag window of ±500 ms. This window duration was selected to be sufficient for capturing multiple theta cycles, thereby enabling the evaluation of rhythmic modulation while minimizing noise from longer timescales. Cell pairs contributing fewer than 10 total spikes were excluded from the analysis.

Given that hippocampal firing sequences are organized specifically according to the trajectory of the animal, we separated spike trains based on movement direction to reveal CCG patterns. A reference vector v_1_ was defined as projecting from the centroid of the place field of cell 1 to the centroid of the place field of cell 2. Movement epochs were classified as running in the direction from cell 1 to cell 2 when the angle between the instantaneous velocity vector of the animal v_2_ and the reference vector v_1_ fell between -90° and 90°. Epochs falling outside this range were classified as the opposite direction.

### Theta index for cofiring

We utilized a theta index to quantify the extent to which the cofiring of cell pairs was modulated by theta oscillations. As the gradual rise and fall of firing rates as an animal traverses a place field introduces a prominent low-frequency component into the CCG, we applied a baseline correction procedure to isolate intrinsic theta oscillatory activity. A low-frequency baseline was estimated by smoothing the raw CCG with a wide Gaussian kernel (window width = 150 ms) and was subsequently subtracted from the raw CCG to obtain the baseline-corrected CCG. We then computed the power spectral density of the baseline-corrected CCG in the 0-25 Hz range. The theta index was defined as the normalized peak power within the theta frequency band, calculated as the maximum power value between 6 and 12 Hz divided by the total integrated power across the 0-25 Hz spectrum.

### Reactivation consistency

LFPs were band-pass filtered (150–250 Hz) to extract ripple-band activity. The instantaneous power was computed using the Hilbert transform and smoothed. SWR events were defined as periods where the ripple power exceeded the mean by at least 3 SD. Events with a duration between 15 ms and 400 ms were retained. SWRs were detected independently on all tetrodes, and the final events were determined by the union of valid time windows across channels.

Reactivation consistency was quantified by assessing the similarity of neuronal co-firing patterns across different SWR events during rest periods (Fig. 6F). For each identified SWR, we constructed a population vector consisting of the spike counts of all recorded place cells during that event. We then computed the pairwise Pearson correlation coefficients between population vectors from all SWR events within a single rest session. The reactivation consistency was calculated as the average of these pairwise correlations, serving as a metric for the stability of the intrinsic neural dynamics during memory consolidation. To ensure robust analysis, only SWR events containing at least 2 active neurons and a total of 3 spikes were included.

### Statistical analysis

All statistical analyses were performed in MATLAB and SPSS. Because multiple observations could be obtained from the same animal, the data had a hierarchical structure, and linear mixed effects models (LMMs) were therefore used for most statistical analyses. In all models, Rat ID was included as a random intercept to account for inter-animal variability and the non-independence of observations from the same animal. For group comparisons that did not involve across-day changes, Group (WT vs. App^NL-G-F^) was included as a fixed effect. For analyses involving changes across days, Group, Day, and the Group × Day interaction were included as fixed effects to assess group differences and their experience-dependent changes. Fixed-effect regression coefficients (β) were reported as estimates of effect size. Statistical significance was set at p < 0.05.

---

**Figure 1–figure supplement 1.**
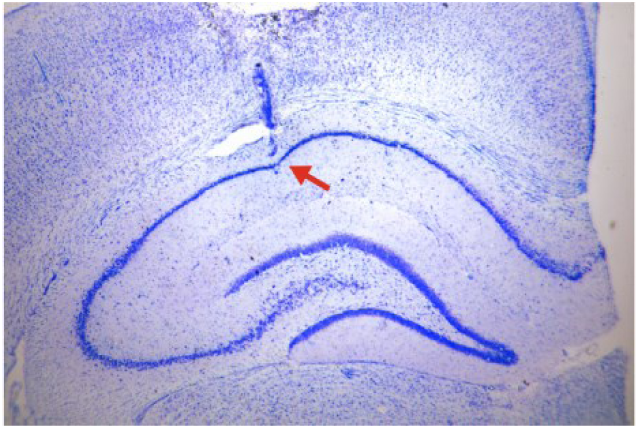
Verification of tetrode locations.

**Figure 1–figure supplement 2.**
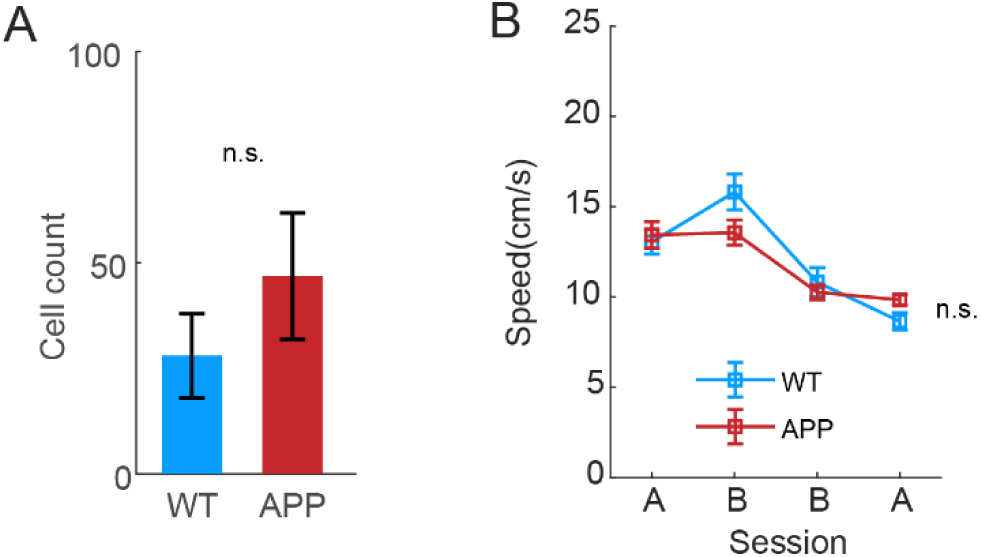
Comparable cell counts and locomotor behavior between WT and App^NL-G-F^ rats. **A.** Number of recorded cells (linear mixed-effects model, F(1, 42) = 1.5, p = 0.2, β = 18.9). **B.** Running speed during open field exploration (F(1, 184) = 0.2, p = 0.7, β = 0.4).

**Figure 1–figure supplement 3.**
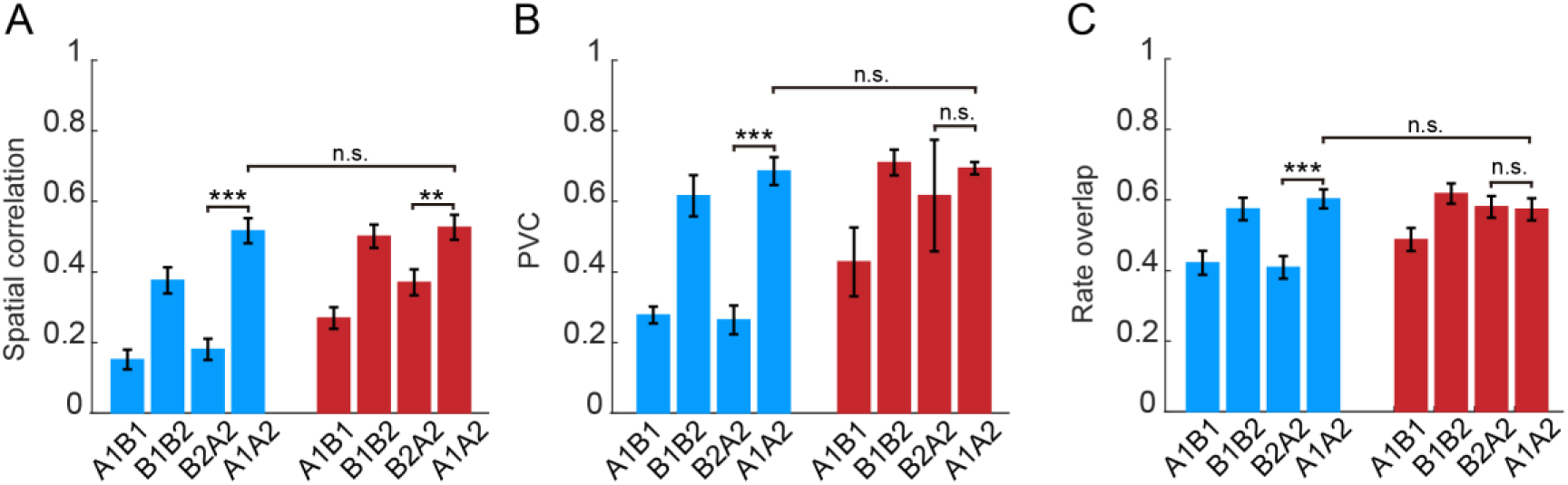
Spatial representation similarity across context transitions. **A**. Spatial correlation across different contexts (linear mixed effects model, group × transition interaction, F(1, 354) = 6.9, p = 0.009; post hoc comparison, A1A2 in two group, t(354) = 0.2, p = 0.8, β = 0.01; WT in A1A2 and B2A2, t(354) = -6.9, p = 2.5×10^-11^, β = -0.3; App in A1A2 and B2A2, t(354) = -3.2, p = 0.001, β = -0.2). **B.** PVC across different contexts (group × transition interaction, F(1, 12) = 7.9, p = 0.02; post hoc comparison, A1A2 in two group, t(12) = 0.09, p = 0.9, β = 0.008; WT in A1A2 and B2A2, t(12) = -5.25, p = 3.8×10^-4^, β = -0.4; App in A1A2 and B2A2, t(12) = -0.8, p = 0.4, β = -0.08). **C.** Rate overlap across different exposures (group × transition interaction, F(1, 341) = 11.0, p = 0.001; post hoc comparison, A1A2 in two group, t(341) = -0.7, p = 0.5, β = -0.03; WT in A1A2 and B2A2, t(341) = -4.5, p = 7.8×10^-6^, β = -0.2; App in A1A2 and B2A2, t(341) = 0.2, p = 0.9, β = 0.007).

**Figure 3–figure supplement 1.**
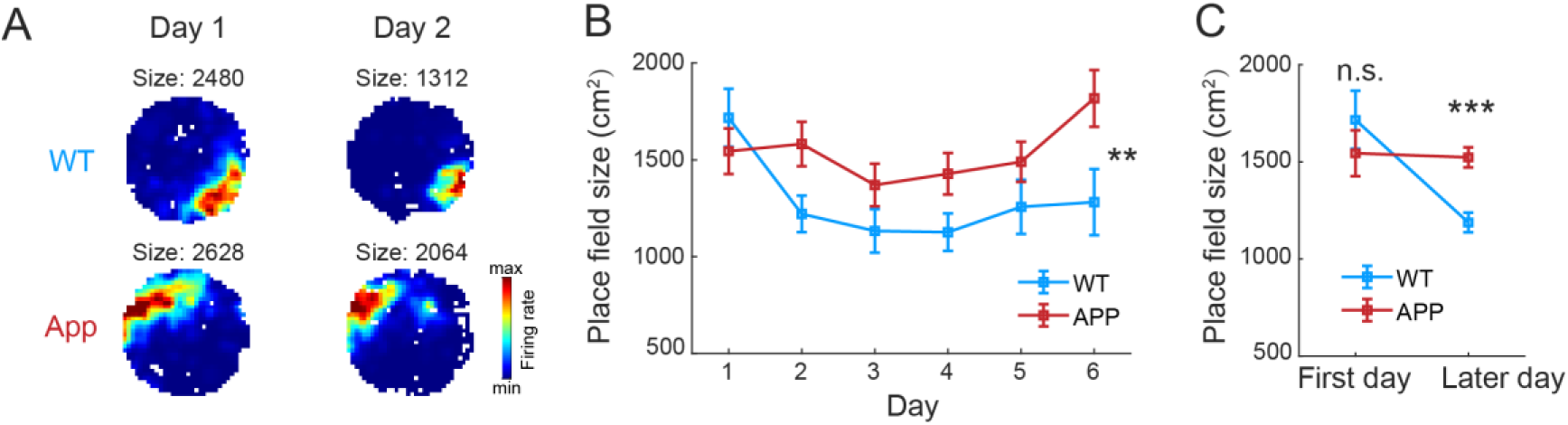
Absence of experience-dependent place field refinement in App^NL-G-F^ rats. **A.** Representative rate maps of a place cell tracked across consecutive days (Day 1 and Day 2) in context B. **B.** Place field size in context B across training days. Data represents the population average of all recorded place cells on each day. **C.** Comparison of place field size in context B between the initial exposure (Day 1) and subsequent training days (aggregated data from Days 2–6).

**Figure 4–figure supplement 1.**
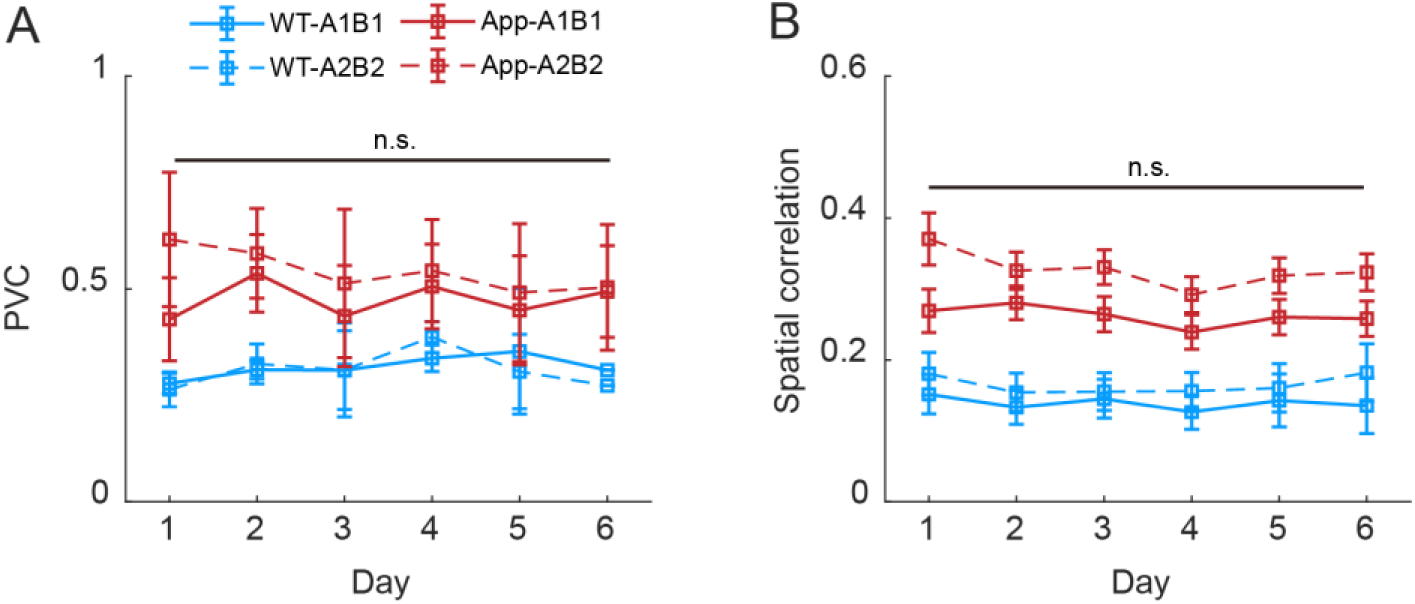
Spatial remapping remains largely unchanged across days. **A.** PVC for A1B1 and A2B2 transitions across days (linear mixed model, main effect of days, t(66) = 0.6, p = 0.6, β = 0.01). B. Spatial correlation for A1B1 and A2B2 transitions across days (t(2680) = -0.7, p = 0.5, β = -0.005).

**Figure 5–figure supplement 1.**
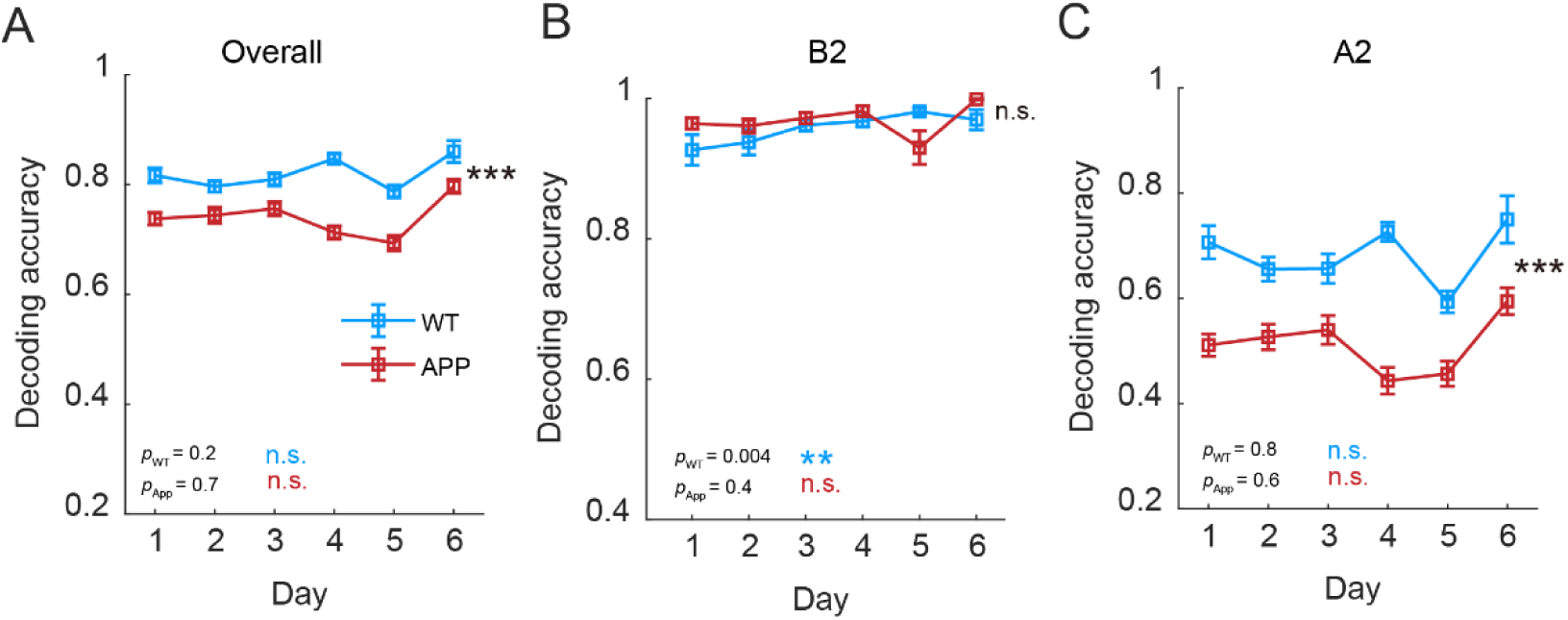
Cofiring based on spatial models shows little significant experience-dependent change in context discrimination. **A.** Overall decoding accuracy (linear mixed effects model, main effect of group, F(1, 491) = 16.6, p = 5.3×10^-5^, β = 0.005. WT: F(1, 223) = 2.0, p = 0.2; App: F(1, 268) = 0.7, p = 0.4). Features are derived from tau based on the expected firing rate. **B.** Accuracy specific to session B2 (main effect of group, F(1, 491) = 3.3, p = 0.07, β = 0.01. WT, F(1, 223) = 8.4, p = 0.004, F(1, 268) = 0.8, p = 0.4). **C.** Accuracy specific to session A2 (main effect of group, F(1, 491) = 27.9, p = 1.9×10^-7^, β = -0.002. WT, F(1, 223) = 0.1, p = 0.8, App, F(1, 268) = 0.3, p = 0.6).

**Figure 5–figure supplement 2.**
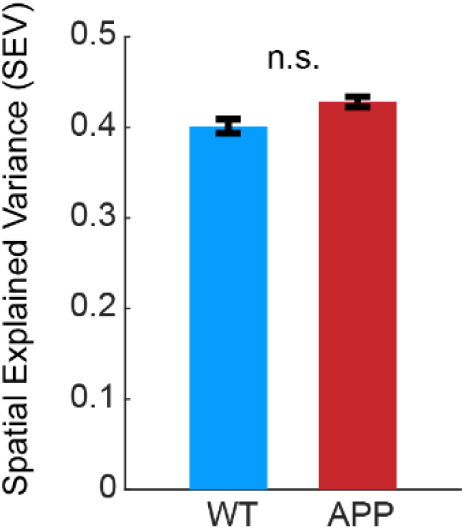
No difference in Spatial Explained Variance between WT and App^NL-G-F^ rats

**Figure 6–figure supplement 1.**
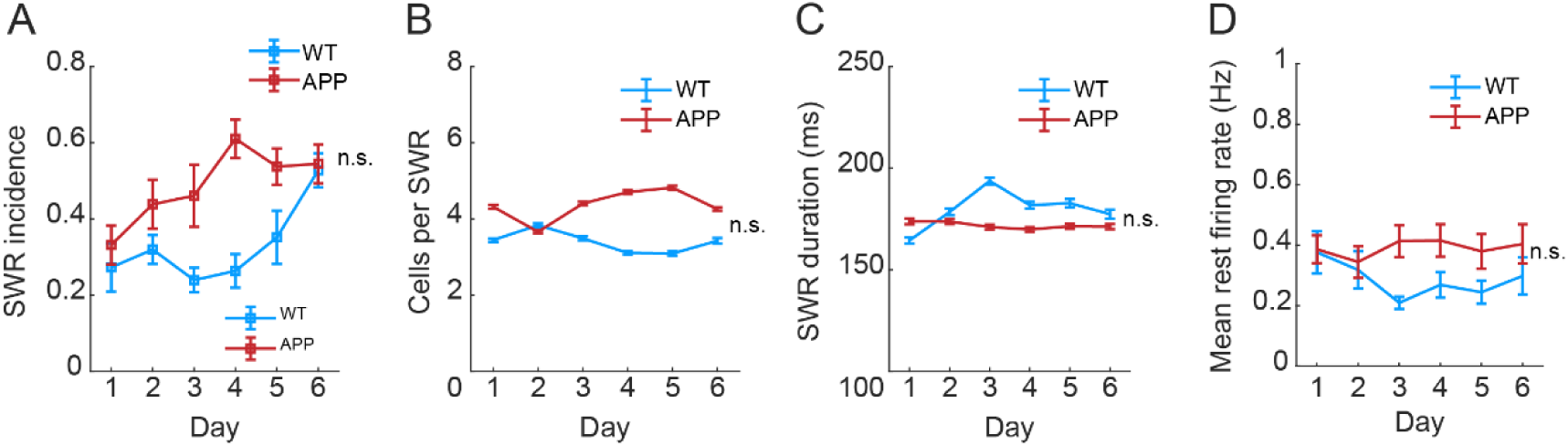
Basic SWR and firing properties remain comparable in WT and App^NL-G-F^ rats. **A.** SWR incidence (linear mixed-effects model, main effect of group, F(1,152) = 0.7, p = 0.4, β = 0.1, n_WT_ = 72 rest epochs in 4 rats, n_App_ = 84 epochs in 4rats). **B.** Number of participating cells per SWR event (F(1,36684) = 0.2, p = 0.7, β = 0.4, n_WT_ =12690 events in 4 rats, n_App_ = 23998 events in 4rats). **C.** SWR duration (F(1,36684) = 0.01, p = 0.9, β = 1.3). **D.** Mean rest firing rate (F(1,1688)=1.5, p=0.2, beta=0.03).

**Supplementary Table 1.** Summary of daily recorded cells.

|  | Rat | Day | Day | Day | Day | Day | Day |
| --- | --- | --- | --- | --- | --- | --- | --- |
| Group | ID | 1 | 2 | 3 | 4 | 5 | 6 |
| WT | 32 | 49 | 55 | 51 | 49 | 56 | 48 |
| WT | 59 | 21 | 29 | 25 | 25 | 17 | 15 |
| WT | 61 | 29 | 28 | 35 | 46 |  |  |
| WT | 65 | 6 | 2 | 5 | 6 | 3 | 2 |
| APP | 22 | 65 | 85 | 69 | 66 | 73 | 72 |
| APP | 23 | 42 | 43 | 46 | 52 | 45 | 40 |
| APP | 34 | 8 | 5 | 5 | 4 |  |  |
| APP | 60 | 47 | 59 | 69 | 76 | 71 | 67 |

